# Evaluating Species at Risk in Data-Limited Fisheries: A Comprehensive Productivity-Susceptibility Analysis of the Most Traded Marine Aquarium Fish

**DOI:** 10.1101/2024.03.26.586872

**Authors:** Gabrielle A. Baillargeon, Alice A. Wynn, Jemelyn Grace P. Baldisimo, Michael F. Tlusty, Andrew Rhyne

## Abstract

The marine aquarium trade is a significant global industry harvesting millions of live coral reef fishes annually. Wild-caught fish dominate public and private aquaria markets in the USA and Europe, supporting fisher livelihoods in the Indo-Pacific. This diverse and species-rich trade is considered data-limited, creating barriers to quantify the sustainability of this fishery to the net benefit of the coral reef socio-ecological system. We present a revised and expanded productivity-susceptibility analysis (PSA) framework to assess the vulnerability to overharvesting of the top 258 traded species, an estimated 92.5% of all import volume into the USA in 2011. Vulnerability was calculated based on various productivity and susceptibility factors, tailored to the unique life-history and fishery selectivity characteristics of the marine aquarium trade. We present novel factors that improve model accuracy, methods to overcome missing data for individual factors, and apply an improved Gaussian mixture model clustering algorithm to objectively classify species as least, moderately, or most vulnerable. Our results show that an overwhelming 85% of species evaluated fall into the least or moderately vulnerable classification, with the remaining species designated as high priority for localized assessment and management initiatives. A comparative case study between our PSA and the popular FishBase Vulnerability assessment illustrates how it is ill-suited to handle data limitations of non-food fishes. The results of our PSA, at a species and family level, provide useful information to stakeholders and serves as a robust and accessible risk assessment tool to prioritize species for management based on their vulnerability scores.

## 1. Introduction

Appropriate fisheries management needs to address the productive capacity of each stock, as well as its susceptibility to overfishing. Without the productivity-susceptibility analysis (PSA), adequate management actions may be challenging to implement. Data limitations often arise given the significant effort required to monitor a large number of species being harvested, along with the multitude of biological, environmental and anthropogenic factors that can influence stock sizes (Hobday *et al*. 2007, McClanahan *et al*. 2023, Baillargeon *et al*. 2020). Fisheries that target marine aquarium trade (MAT) species are recognized for their diversity and often small harvest volumes (Rhyne *et al*. 2017). Due to the high diversity and low volume characteristic of this fishery, a majority of species in the MAT remain wild-caught (Tlusty *et al*. 2013). Although there have been recent technological advancements paving the way for aquaculture to supplement wild-caught fish, there remains a limited number of species where commercial aquaculture is viable (Tlusty, 2002). Commercial aquaculture is concentrated in western countries who have historically played the role of major importing nations in the trade, instead of supplementing fishing livelihoods with cultured production in native ranges (Tlusty, 2002).

Species harvested for the MAT are particularly data-deficient as they are: (1) less voluminous in trade compared to food fish, (2) often harvested from remote, biodiversity hotspots throughout the coral triangle, (3) and remote locations limits catch reporting in a traditional framework that results in long term datasets (Wood 2001a; Wood 2001b; Fujita *et al*. 2014; Dee *et al*. 2014; Okemwa *et al*. 2016; Dee *et al*. 2019; Baillargeon *et al*. 2020; Biondo and Burki 2020). A successful method to rapidly assess the vulnerability of a species to fishing activity within a data-limited context is the productivity-susceptibility analysis (Hobday *et al*. 2007, Patrick *et al*. 2009). The PSA estimates a stock’s productivity based on widely known life history traits that are tailored to key growth indicators and trophic niches specific to a group of fishes. To assess fishery and trade influences on a stock’s abundance, the PSA also scores the susceptibility of a fish to fishing pressure across several factors, and productivity and susceptibility are combined into a final vulnerability score (Hobday *et al*. 2007, Hobday *et al*. 2011). The MAT PSA developed by Baillargeon et al. (2020) focused on 32 species, comprising the top 20 species in trade along with 12 species assessed by other PSA studies. Seven productivity and five susceptibility factors were deemed robust across all species and were used to calculate vulnerability (Baillargeon *et al*. 2020).

Rhyne *et al*. (2012, 2017) reported that the United States is the largest importer of marine aquarium fish, having imported approximately 2,300 species from over 40 countries within a four-year period. Similar trends in imports are observed in the UK and the European Union (Rhyne *et al*. 2017). However, it’s notable that only a limited number of these species have undergone assessment or are covered by active management plans to evaluate their vulnerability to overfishing. The value chain for marine aquarium fish is markedly different from that of food fisheries. These species are captured, transported, sold, and maintained alive, significantly increasing their economic value. The value typically escalates towards the end of the supply chain, where the price of a fish can substantially increase by the time it reaches the supplier in the importing country. Within a given genus, the value of species varies considerably. For instance, juvenile or deep-water species, which are often challenging to capture, and visually appealing or colorful species, tend to be more sought after by consumers, further influencing their market value (Wood 2001a; Bruckner 2005; Rhyne *et al*. 2017).

In 2019, the CITES parties agreed to conduct a review and technical workshop to help understand trends in the MAT and identify species at risk of overexploitation, highlighting the need for a robust and accessible assessment method for managers to implement at the local to global scale (CoP19 Inf. 99). The ease and accessibility of FishBase’s (Froese and Pauly, 2023) vulnerability calculation has recently been highlighted on an international scale, as it is the leading species-specific assessment method supporting the most recent CITES report (UNEP-WCMC, 2022) evaluating the global MAT. However, the report lacks transparency of methods and data. FishBase’s current model has limitations as a resource for fishery vulnerability information across species, where data may be missing (Cheung *et al.,* 2005). FishBase vulnerability scores have been shown to be ineffective at modeling risk for reef fishes on a global scale and when handling species which do not conform to growth patterns the model is based on (Go *et al*. 2015). Therefore, although it is the leading analysis metric referenced it may not be the most useful in a management context.

To ensure appropriate resource extraction of the MAT species and sustainability of the MAT, it is critical to perform rapid fisheries assessments that provide accurate results and do not require extensive fisheries management resources. Therefore, we expanded the PSA model developed by Baillargeon *et al*. (2020) to 258 species, which represent approximately 92.5% of individuals imported into the US by count (Rhyne *et al*. 2015). We addressed the issue of data deficiency by implementing methods to overcome data gaps, quantified factors that are most impactful to a fish’s vulnerability across the diverse 258 species and improved the specificity of the PSA framework and vulnerability characterization to marine aquarium fish.

## 2. Materials & Methods

### 2.1 Assessing variables to include in PSA

Identifying the optimal set of factors to obtain the vulnerability score and mitigate uncertainties determines the effectiveness of a PSA framework. To achieve this, the PSA model developed by Baillargeon *et al*. (2020) was applied to the top 258 MAT species imported into the US based on 2011 data (Rhyne et al. 2015, 2017). The feasibility and practicality of use was evaluated assessing data availability in each productivity and susceptibility factor being scored. When species level data was unavailable, a congener, closest genetic relative, or family level data was used. In the absence of data, experts on reproductive biology of marine ornamental fish species were consulted to provide fecundity data needed for this study. Model sensitivity was also tested by looking at the influence of each productivity and susceptibility factor on the resulting vulnerability score. Validation of the method developed by Baillargeon *et al*. (2020) resulted in changes to the productivity and susceptibility factors being assessed and the weighting for each factor (see Table S1 for complete data binning and scoring matrix).

Productivity and susceptibility factors were scored for each species individually on a 1−3 (low−high) scale, in line with Patrick *et al*. (2009) and Hobday *et al*. (2011) (see Table S1 for scoring matrix). Productivity is an indirect measurement of a species’ ability to reproduce and indicates resiliency to changing environmental conditions (Baillargeon *et al*., 2020). Five productivity factors: maximum size, mean trophic level, breeding strategy, fecundity, and pelagic larval duration (PLD) were included in this PSA framework (Table 1).

**Table 1:** Productivity and susceptibility factors with corresponding data sources. Bold text indicates a factor weight of 2 in the model.

| Factor | Data sources |
| --- | --- |
| **Productivity** |  |
| **Maximum size** | Primary literature, Michael, S. (2005), FishBase |
| Mean trophic level | FishBase, Primary literature |
| Breeding strategy | FishBase, Breder and Rosen (1966), Thresher (1984) |
| **Fecundity** | Primary literature, Hobbyist forums (i.e.: breedersregistry.org, mbisite.org), FishBase |
| Pelagic larval duration | Primary literature |
| **Susceptibility** |  |
| **Ecological niche/distribution** | Eschmeyer's Catalog of Fishes, IUCN Red List assessments, FishBase |
| Cyanide use | Aquariumtradedata.org |
| Encounterability depth | IUCN Red List assessments |
| Aquarium suitability | Online hobbyist databases, (LiveAquaria.com, Saltcorner.com, Reeflex.net), Michael, S. (2005) |
| **Trade volume** | Aquariumtradedata.org |
| **Life cycle stage of harvest** | LiveAquaria.com, BlueZooAquatics.com |

Susceptibility measures the likelihood that fishing pressures will have a negative impact on a species’ population (Patrick *et al*. 2009). Susceptibility has a reverse scale from productivity, where high susceptibility translates to a higher vulnerability score. Six susceptibility factors: geographic distribution, encounterability depth, suitability for aquarium keeping, volume in trade, and the life stages harvested, and cyanide were included in this PSA framework (Table 1).

Validity of scientific names for species included in this study were checked against the online version of Eschmeyer’s Catalog of Fishes up to December 2023 (Fricke *et al*. 2023). Data for the top 258 traded species were sourced from a mix of primary literature, open-source databases and repositories, and aquarium hobbyists’ gray literature (Table 1). An analysis of change in vulnerability scores across the 32 species analyzed in Baillargeon *et al*., 2020 utilizing the present model was also conducted (Figure S1).

A phylogenetic tree with vulnerability score heatmap data was created for the 258 species list using the *rtol*, *ape*, and *ggtree* packages in R (Figure S2). Organizing the data phylogenetically helps to visualize vulnerability score trends across taxonomic levels. Phylogenetic data was sourced from the Open Tree of Life database.

### 2.2 PSA Mathematical Framework

Productivity was calculated based on six life history factors (Table 1). A weighted arithmetic mean was used to calculate Productivity, where is the productivity score and is the factor weight. Increasing the factor weight to 2 represents the factor’s importance in determining the vulnerability of a species in a fishery.

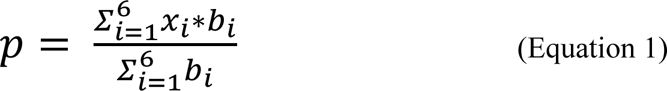

Susceptibility was calculated by using a weighted mean of logarithms expressed as an exponential function, with base 10 raised to the power of the weighted logarithmic mean, where *y_i_* is the susceptibility factor score and a*i* is the factor weight:

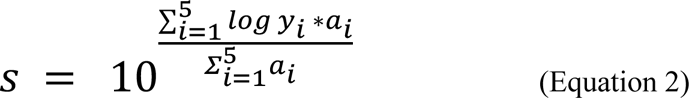

These factors were quantified in a data-binning process where productivity and susceptibility scores are calculated separately then inputted into the Euclidean distance formula to output the vulnerability score (*v*). Where *p* is the mean productivity score, *s* is the mean susceptibility score, and *v* is the vulnerability score (equation 3). Vulnerability is functionally the distance from the origin (1,3) of the productivity-susceptibility plot.

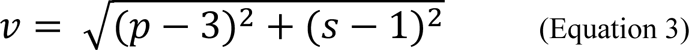

### 2.3 Productivity and Susceptibility Factor Refinement

#### 2.3.1. Productivity factors

Length at maturity and maximum age factors were removed in this analysis due to lack of reliable data for a majority of the 258 species being assessed. In addition, when the impact of each factor was examined, maximum size was closely correlated to the maximum age and length at maturity (R^2^>0.9). Maximum size was given double-weight among the productivity factors to account for its representation of numerous factors.

The factor breeding strategy, was adjusted to be dependent on fecundity in cases of high disparity between breeding strategy and fecundity values to ensure accuracy (Table S2). Because of this scaling, score weight for this factor was set to 1 instead of 2 to avoid double-counting within the model. All other factors and scoring bins remained unchanged from Baillargeon *et al*. 2020.

#### 2.3.2 Susceptibility factors

Ecological niche and geographic distribution were combined into a single factor in this model, to account for the interaction of species range and habitat specificity impacting overall susceptibility to fishing effort. Geographic distribution (categorized as large or small) was determined by referring to published IUCN Red List Assessments (IUCN 2022) and Eschmeyer’s Catalog of Fishes (Fricke *et al*. 2023). This was then cross-referenced with habitat specificity information from FishBase (Froese and Pauly, 2023) which was categorized as wide or narrow. Data from ecological niche and geographic distribution were then binned into a single-factor score following Rabinowitz (1981) methodology (Figure 1). For example, species with small geographic ranges and narrow habitat specificity were classified as most susceptible, with a score of 3.

**Figure 1:**
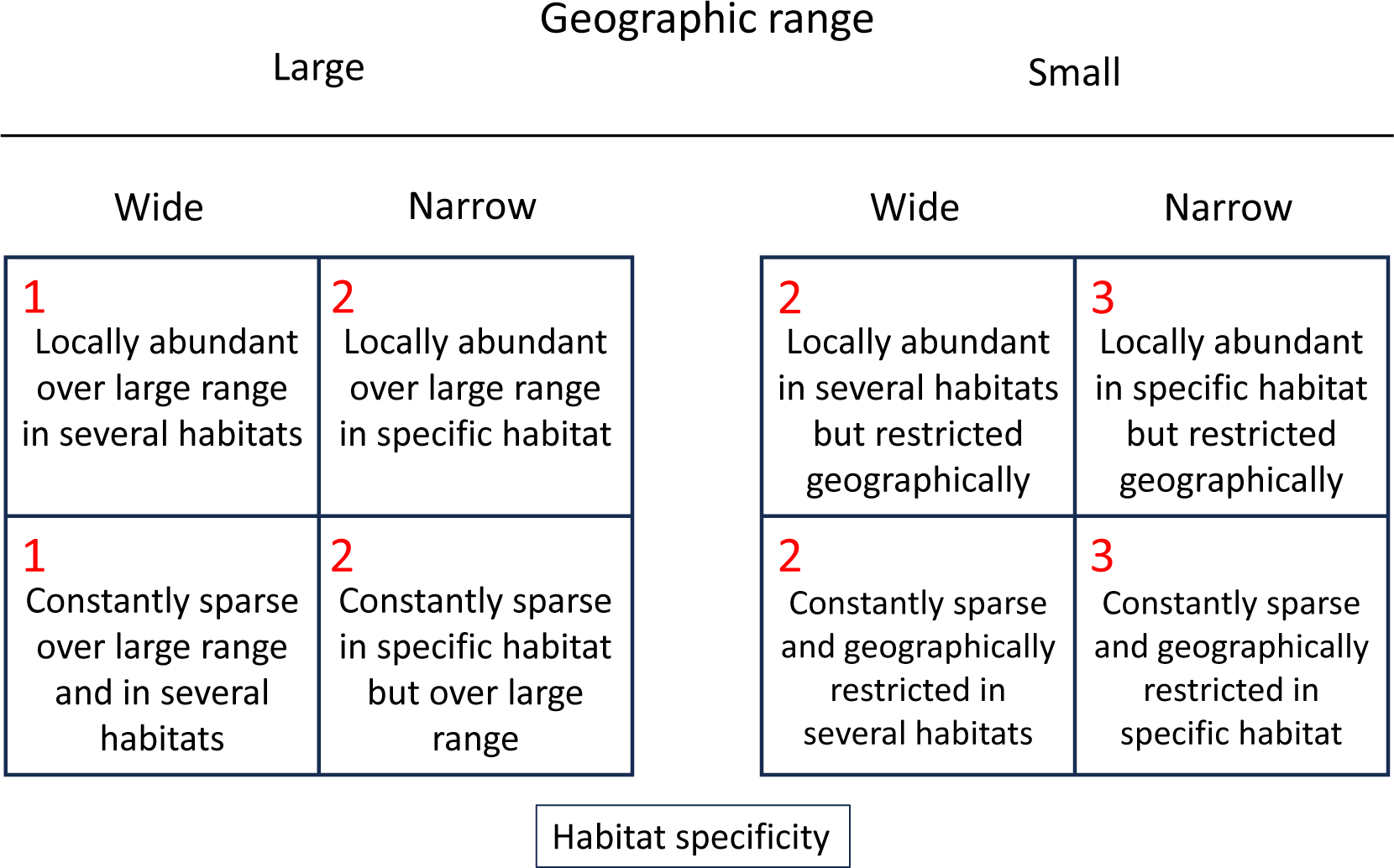
Scoring matrix for susceptibility factor Ecological niche/distribution based on Seven Forms of Rarity (Rabinowitz, 1981). Ecological niche/distribution scores were calculated based on their geographic range determined by the California Academy of Science and IUCN databases, combined with habitat specificity from FishBase. Factor score is indicated in the upper left corner of each categorization bin.

Encounterability depth was obtained from the depth range of each species indicated in the IUCN Red List database (IUCN 2022) and scored based on the maximum depth, instead of the average depth (Baillargeon *et al*., 2020) where a species can be observed.

Aquarium suitability was scored by converting descriptive care levels estimated by hobbyist sources (Table 1) into a 5-point numerical scale (least difficult = 1 to most difficult = 5). These hobbyist databases provided various phrases used to describe care difficulty. Terms such as “Easy” and “Beginner” represented the low points of the scale (least challenging). Phrases such as “Moderate,” “Average,” and “Intermediate” represented the middle points of the scale (intermediate). Lastly, the highest (most difficult) part of the scale was represented by phrases such as “Expert,” “Advanced,” “Professional,” and “Not suitable for home aquaria.” In the majority of cases, multiple hobbyist databases had information on a particular species, and care level was averaged across these sources. This 5-point scale was converted to the 3-category PSA framework by assigning 1 for scores <2.5 (Less difficult), 2 for scores between 2.5 to 3.5 (Moderate difficulty), and 3 for scores >3.5 (More difficult). This gives a quantifiable range to narrative data, in comparison to the “easy, medium, hard” categorization from the same data sources in Baillargeon *et al*. 2020.

When fishes are harvested at a rate faster than their reproduction rate, overexploitation occurs (Aubone, 2004). Thus, species with a high volume in trade and high fecundity may be considered less susceptible to overexploitation compared to species with low to moderate volume in trade and low fecundity. To capture this in the revised PSA model, the factor score for trade volume was scaled based on the species’ productivity score. First, annual trade volume was binned into three categories: <3,000 individuals; 3,000 to 15,000 individuals; and >15,000 individuals. Data bins are based on the 1^st^ and 3^rd^ quartiles from the spread of volume in trade data. Since productivity scores ranged from 2.30 to 0.77 in this analysis, if a species was highly productive (*i.e.* 2.30 to 2.00), trade volume was considered low and scored as a 1, regardless of the import volume. Species with moderate productivity (*i.e.* 2.00 to 1.90) and intermediate to high trade volume (3,000 to >15,000 individuals) were scored 2, while species with low trade volume or <3,000 were scored 1. Finally, species with low productivity (1.90 to 0.77) and moderate to high import volume (3,000 to >15,000) were scored 3, or at 2 if the number of species traded was <3,000 at this productivity level. This factor was weighted at two due to the importance of trade volume on fishery vulnerability (Table 1, Table S1).

The life cycle stage at harvest (LCSH) was added as a susceptibility factor due to its impact on local population dynamics. Harvesting adults negatively impacts the reproductive potential of a stock, including its ability to withstand and recover from fishing pressure (Begg and Marteinsdottir 2003; Law 2007). Fishing juveniles may reduce the population of breeding adults in the future and affect population renewal (Wood 2001a). As such, species harvested in both juvenile and adult stages were considered more susceptible to overfishing (score of 3) than species harvested only as juveniles (score of 1) or adults (score of 2). This scoring framework was developed based on the idea that removing all life stages significantly reduces potential recruitment, as does removing adult broodstock, whereas removing non-breeding juveniles has less direct impact on population size (Table S3). LCSH on a per species basis was confirmed by referencing size classes of fish sold by well-known online retailers (Liveaquaria.com, Bluezooaquatics.com). In the absence of available data, an expert in aquarium fish rearing and trade was consulted.

### 2.4 Sensitivity Analysis and Model Comparison

A model sensitivity analysis was done to test the impacts of individual factor scores, weighting, and combined effects across multiple factors for both productivity and susceptibility. Multiple factor scores were manipulated, individually and then in succession, while all other factors were set at a neutral score of 2 (Figure S3). For productivity, the following grouped factor scores were independently manipulated from 1 (low) to 3 (high): fecundity (weighted), breeding strategy, and pelagic larval duration. Similarly, a group of susceptibility factors (aquarium suitability, LCSH, and encounterability depth) were manipulated from 3 (high) to 1 (low). This analysis compared the change in vulnerability score across three weighted and unweighted factors. This three-factor manipulation included an equal number of weighted (1) and unweighted factors (2) across productivity and susceptibility factors. An expanded model sensitivity analysis was conducted to reveal trends across single weighted and unweighted factors, with all other factors set at a neutral score of 2 (Table S4). Single productivity factors manipulated include maximum size and trophic level, while susceptibility factors manipulated include ecological niche + distribution and aquarium suitability.

PSA species outcomes were directly compared with the widely available and open-access repositories and assessments for such a large volume of species: IUCN Red List and FishBase. The conservation status of the 258 species in this study was retrieved from the IUCN Red List website (IUCN 2022) and categorically compared to sustainability categorization of PSA vulnerability scores based on our clustering methods. Vulnerability scores were scraped from FishBase.org for all 258 species using R.

### 2.5 Gaussian Mixture Modeling to Classify Species Vulnerability

A semi-supervised machine learning algorithm was implemented to classify all 258 species evaluated into three distinct vulnerability clusters: “most vulnerable”, “moderately vulnerable”, and “least vulnerable”. Multivariate, finite Gaussian mixture models and K-means clustering algorithms were evaluated for best model fit using the open-source *R Mclust* package and *stats* package, respectively. K-means clustering groups species into circular shaped clusters where the borders are defined by minimizing the distance from the centroid center to radius edge based on two-coordinate points. The GMM model was chosen over the K-means for its ability to cluster points along 3 axes based on productivity, susceptibility, and vulnerability values with unequal volume and shape (Table S5).

The silhouette coefficient was evaluated to determine the optimal number of clusters for the expanded dataset. Log-likelihood and Akaike Information Criterion (AIC) model comparison led to the selection of the Mclust VVI finite GMM model. The main distinction between this new GMM and the former GMM used in the previous study (Baillargeon *et al*. 2020) is clustering based on productivity, susceptibility, and vulnerability scores instead of only along the x-y axis. Further, this GMM model framework is set to cluster diagonally from the origin while allowing the volume and shape of each cluster ellipse to differ instead of holding size and shape constant like k-means or other GMM clustering models. Full model code is available here: https://github.com/gbaillargeon/PSA-Gaussian-Mixture-Model.git.

## 3. Results

Across the 258 species, productivity scores ranged from 1.29–3.00, and susceptibility scores ranged from 1.00–2.32, resulting in a vulnerability range of 0.19–1.82 (Table 2). Significant variation in average factor scores between high and low vulnerability scoring species was observed (ANOVA, p<0.05). For productivity factors, maximum size and fecundity represented the largest difference in average values of fish in the top 10 most and least vulnerable clusters (Figure 2).

**Figure 2:**
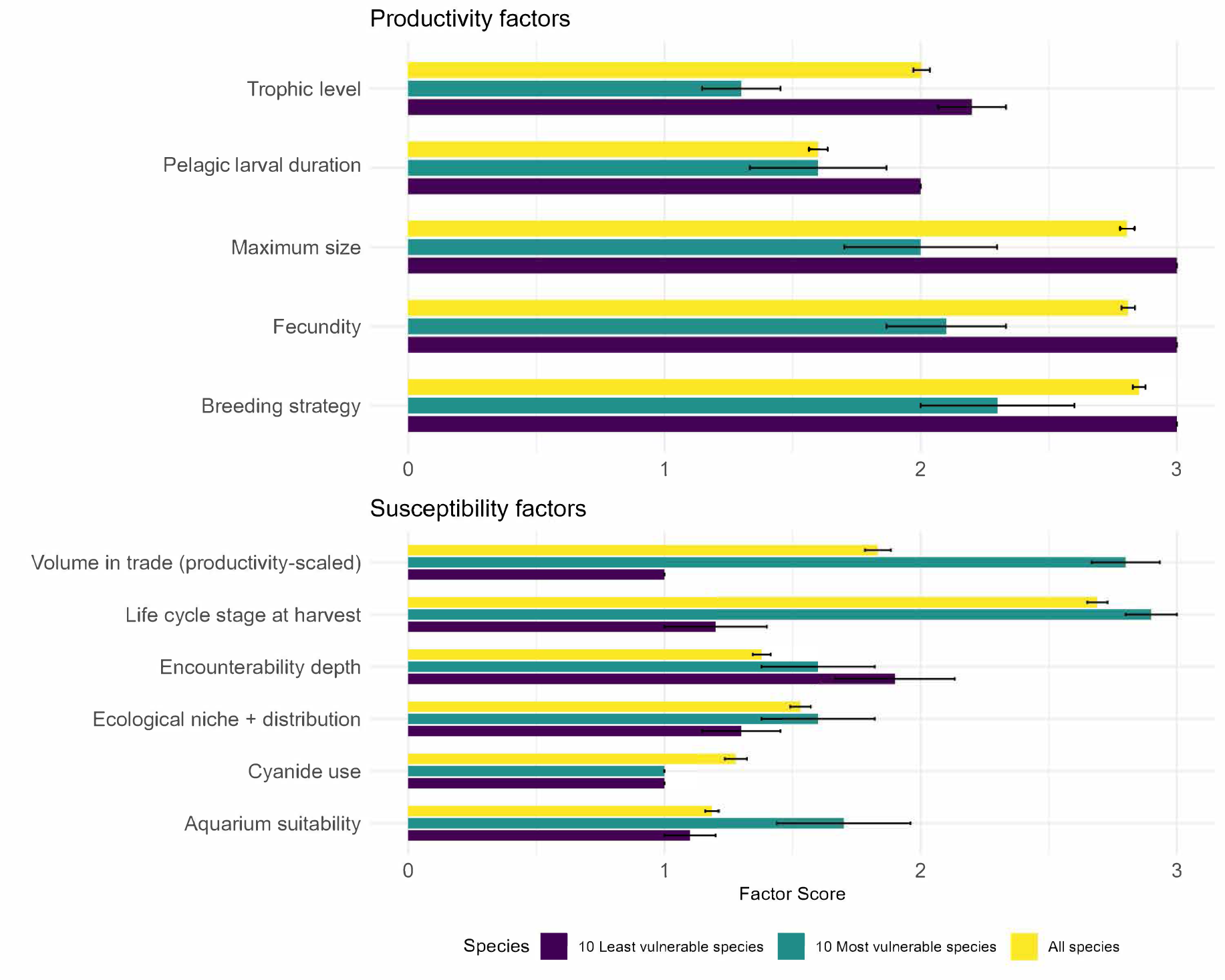
Comparative analysis of average productivity and susceptibility factor scores. Average factor scores (± standard deviation) across five productivity and six susceptibility factors for most vulnerable species (n=10), least vulnerable species (n=10), and all species (n=258).

**Table 2:** Species, Family, Productivity (*P*), Susceptibility (*S*), Vulnerability (*V*), Rank in Trade, IUCN status, Availability as captive-bred for 10 of the most and least vulnerable species assessed. Rank in trade of 1 represents the most imported fish into the US in 2011 (Rhyne et al. 2015). Captive bred availability of species is based on Pouil et al. 2020 with the categories: has been successfully bred in captivity, scare, moderate, or common availability. IUCN status in descending order of vulnerability: Endangered (EN), Vulnerable (VU), Least <u>concern (LC), and Not Threatened (NT).</u>

| Species | Family | P | S | V | Rank<br>in<br>trade | IUCN<br>Status | Availability as<br>captive-bred (Pouil<br>et al. 2020) |
| --- | --- | --- | --- | --- | --- | --- | --- |
| Top 10 most vulnerable species |  |  |  |  |  |  |  |
| *Chiloscyllium punctatum* | Hemiscyllidae | 1.29 | 1.61 | 1.82 | 251 | NT | Moderate |
| *Echidna nebulosa* | Muraenidae | 1.57 | 1.63 | 1.56 | 70 | LC | - |
| *Pterapogon kauderni* | Apogonidae | 2.00 | 2.15 | 1.52 | 9 | EN | Common |
| *Rhinomuraena quaesita* | Muraenidae | 1.57 | 1.49 | 1.51 | 277 | LC | - |
| *Gomphosus varius* | Labridae | 2.00 | 2.05 | 1.45 | 168 | LC | - |
| *Exallias brevis* | Blenniidae | 2.43 | 2.32 | 1.44 | 95 | LC | - |
| *Ostorhinchus parvulus* | Apogonidae | 2.43 | 2.32 | 1.44 | 170 | LC | - |
| *Diodon hystrix* | Diodontidae | 1.86 | 1.76 | 1.37 | 105 | LC | - |
| *Diodon holocanthus* | Diodontidae | 1.86 | 1.76 | 1.37 | 160 | LC | - |
| *Zoramia leptacanthus* | Apogonidae | 2.14 | 2.05 | 1.36 | 104 | LC | Scarce |
| **Top 10 least vulnerable species** |  |  |  |  |  |  |  |
| *Chrysiptera unimaculata* | Pomacentridae | 2.86 | 1.13 | 0.19 | 203 | LC | - |
| *Chromis opercularis* | Pomacentridae | 2.71 | 1.00 | 0.29 | 176 | LC | - |
| *Amblypomacentrus breviceps* | Pomacentridae | 2.71 | 1.00 | 0.29 | 224 | LC | - |
| *Neoglyphidodon nigroris* | Pomacentridae | 2.71 | 1.08 | 0.30 | 37 | LC | Has been bred in captivity |
| *Pomacentrus amboinensis* | Pomacentridae | 2.86 | 1.28 | 0.31 | 132 | LC | Has been bred in captivity |
| *Neoglyphidodon oxyodon* | Pomacentridae | 2.71 | 1.13 | 0.31 | 49 | LC | - |
| *Dischistodus prosopotaenia* | Pomacentridae | 2.71 | 1.17 | 0.33 | 261 | LC | - |
| *Neoglyphidodon melas* | Pomacentridae | 2.71 | 1.26 | 0.39 | 33 | LC | Scarce |
| *Amblyglyphidodon curacao* | Pomacentridae | 2.71 | 1.26 | 0.39 | 92 | LC | Has been bred in captivity |
| *Amblyglyphidodon ternatensis* | Pomacentridae | 2.71 | 1.26 | 0.39 | 107 | VU | Has been bred in captivity |

For susceptibility factors, volume in trade and life stage at harvest showed the greatest difference between top 10 most and least vulnerable species (Figure 2). Ecological niche + distribution, cyanide use, and pelagic larval duration had the least change in average score when comparing the highest and lowest scoring species in terms of vulnerability, indicating these factors had the least influence on the model output (Figure 2). If we assess the most traded species by looking at the top 25% of species in the trade by import quantity, this corresponds to volume in trade rank 1-64 where 15,000 individuals or more are traded annually. Within this commonly traded group, 57.8% of species fall into the ‘least vulnerable’ category, with three of the top ten lowest scoring species (*Neoglyphidodon nigroris, Neoglyphidodon oxyodon* and *Neoglyphidodon melas*) in the PSA being represented. Only four ‘most vulnerable’ species are traded at this volume, accounting for about 10% of the total species considered most vulnerable by our PSA.

Every factor evaluated in this PSA had data available for all species, except for Breeding Strategy, Pelagic Larval Duration, and Fecundity. For breeding strategy 75% of fish evaluated have species level data, pelagic larval duration has 51.9%, and fecundity only have 27% species-specific data (Figure 3). Fecundity has the largest combination of all data types, where breeding strategy was largely determined by species and family level estimates exclusively.

**Figure 3:**
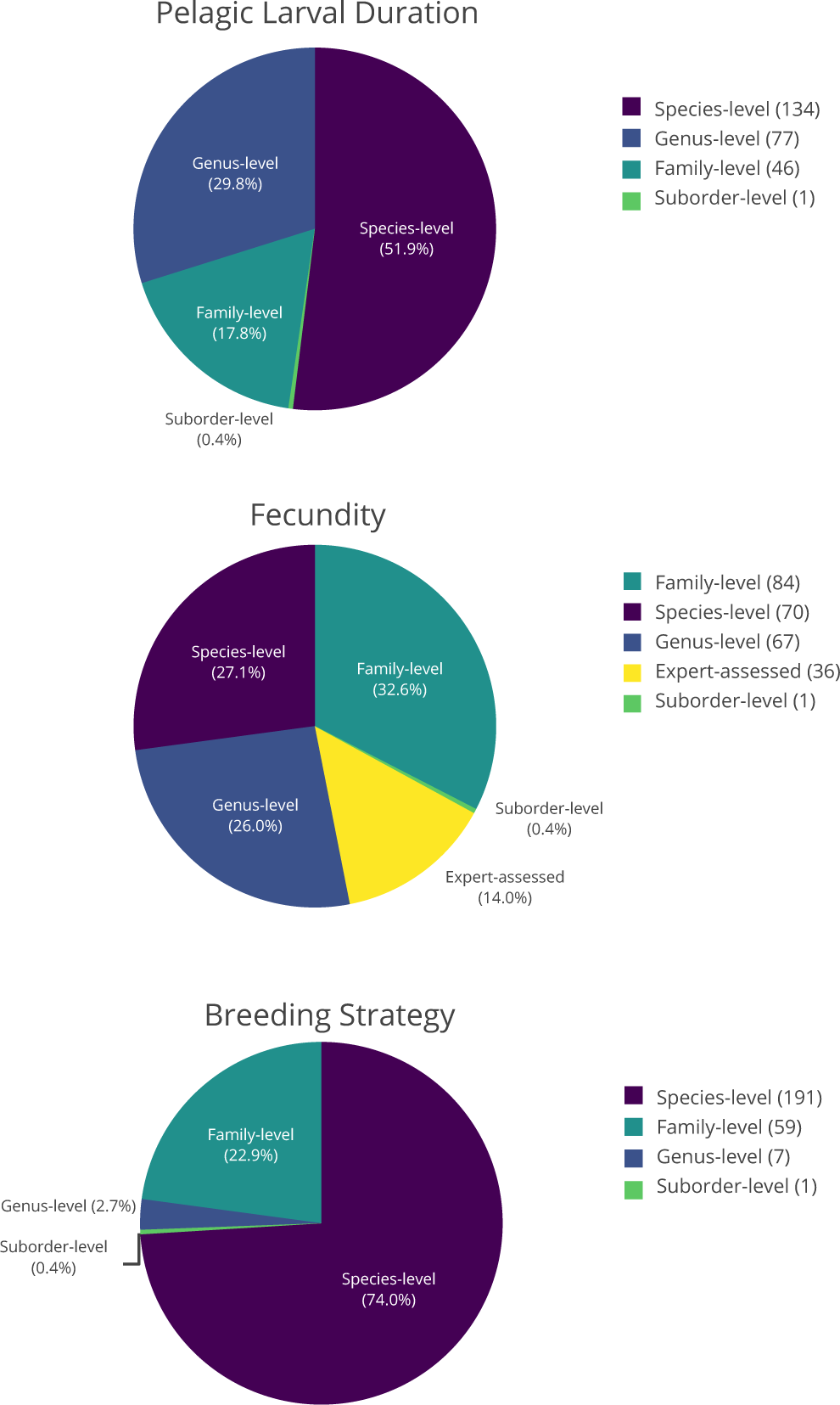
Percentage breakdown for species level data availability for data-limited productivity factors (pelagic larval duration, fecundity, and breeding strategy). For all species assessed (n=258), data for factor scoring was drawn from either the species, family, genus, or suborder level based on data availability for each species. For the factor fecundity, experts on reproductive biology of marine ornamental fish species were consulted to when data was missing at the species or family level. This is displayed within the pie chart (%) with the number of individuals per category demarcated in the legend of each factor.

**Figure 4:**
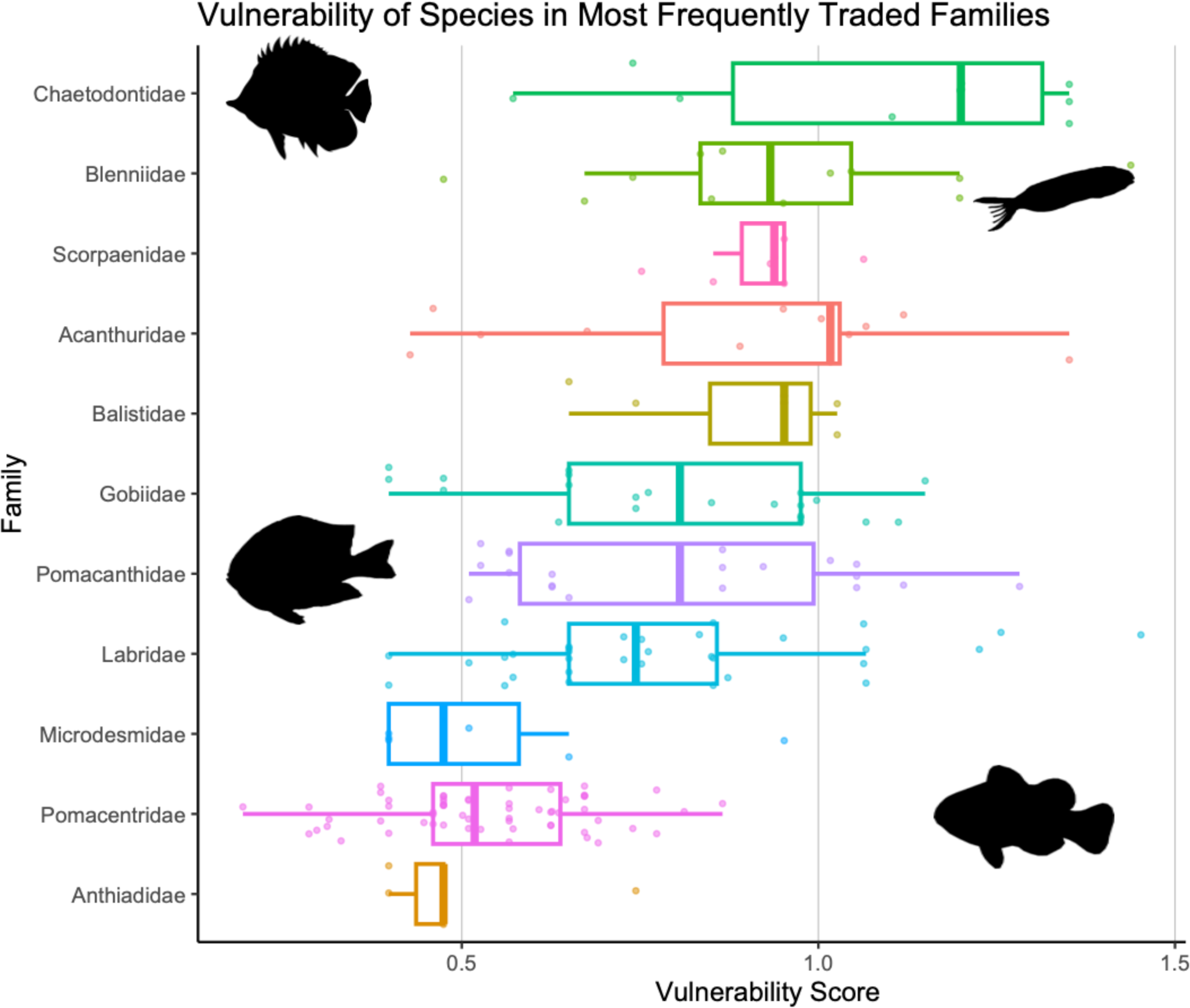
Boxplot of vulnerability scores for the top families by volume in the trade (n=11), ordered from most to least vulnerable. A total of 36 families were assessed, and one-third of those families contained only a single traded species. Notably, 7 families (22% of all families evaluated), account for 69% of all species evaluated in this study. The damselfishes (Family: *Pomacentridae*), the largest family of fish in the analysis, have the lowest average vulnerability of the top 7 families (*v* = 0.53 ± 0.14, n=60), with the three least vulnerable fish (*Chrysiptera unimaculata*, *Chromis opercularis,* and *Amblypomacentrus breviceps*) in the analysis coming from this family (*v* = 0.39). The butterflyfishes (Family: *Chaetodontidae*) are the most vulnerable of these families (*v* = 1.08 ± 0.28, n=10) (Figure 4).

The model sensitivity analysis highlights that productivity has a greater effect on the vulnerability score when manipulating a group of factors as opposed to susceptibility. All factors aside from those being manipulated were set at a “neutral score” of 2. All scores set to neutral produce a vulnerability score of 1.55 for the example species. Shifting a group of productivity factors (fecundity, breeding strategy, and PLD) from 1 to 3 results in a vulnerability score decrease of 0.88, while a scoring shift of 3 to 1 for a group of susceptibility factors (aquarium suitability, encounterability depth, and LCSH) results in a 0.73 decrease in vulnerability score (Figure 5).

**Figure 5:**
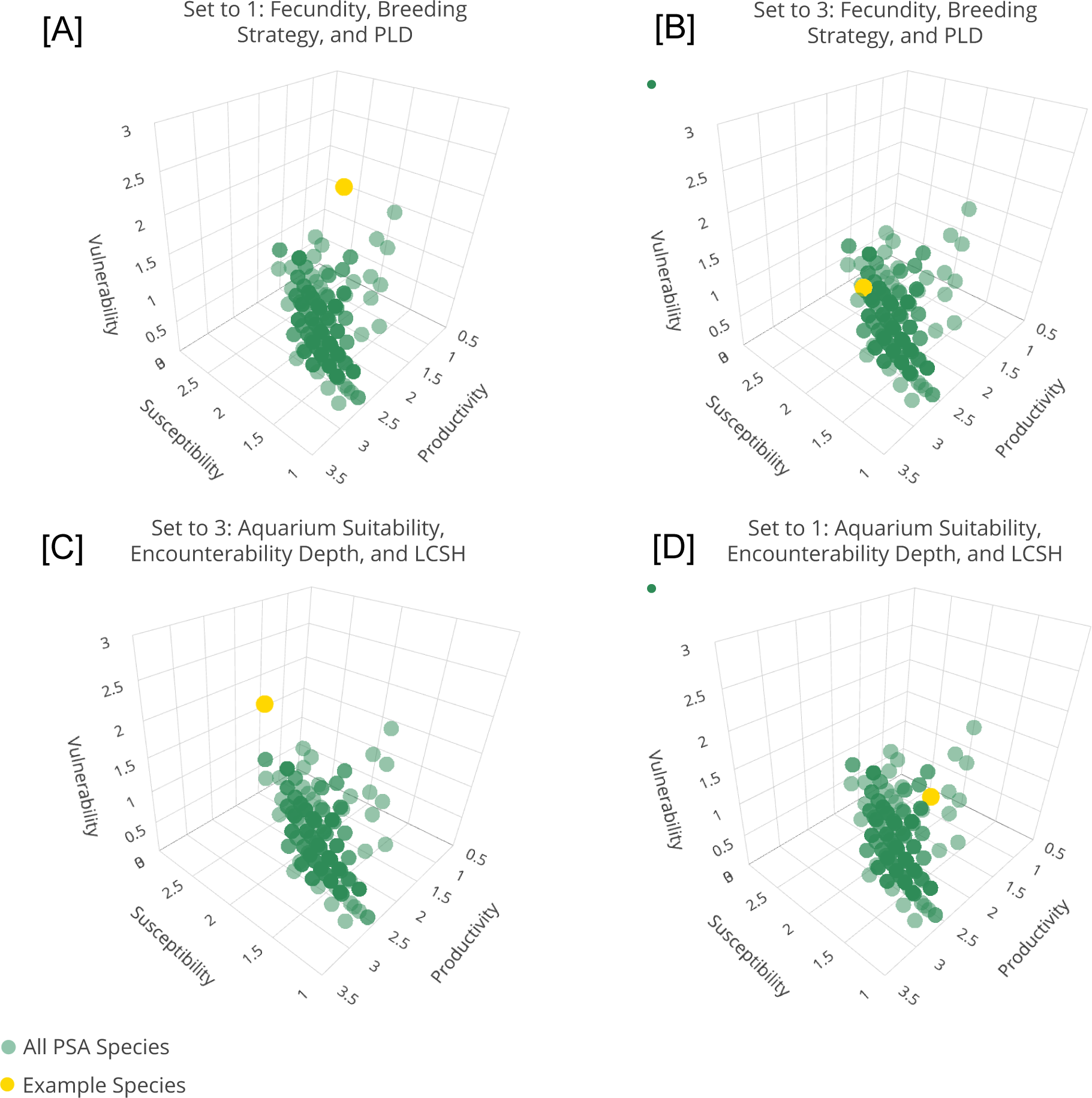
Model sensitivity analysis showing the impact of changing factor scores on the vulnerability score. Species in this analysis are represented by green points (n=258), and the hypothetical example fish is represented by a yellow point. [A] Shows a hypothetical fish with a low productivity score, falling into the most vulnerable category [B] Shows a hypothetical fish with high productivity having a much lower vulnerability score. [C] Shows a hypothetical fish having a similarly high vulnerability score as [A] in the most vulnerable category but due to a high susceptibility score, whereas [D] shows that a low susceptibility score translates to a low vulnerability score, similar to [B].

In the expanded sensitivity analysis, manipulation of single factors revealed a marginal tendency for susceptibility to have a greater effect on the vulnerability score opposed to productivity. Shifting a single weighted productivity factor (maximum size) from 1 to 3, and weighted susceptibility factor (ecological niche + distribution) from 3 to 1 resulted in a vulnerability score decrease of 0.36 and 0.39, respectively. Shifting of single unweighted productivity and susceptibility factors (trophic level and aquarium suitability) using the same parameters resulted in a vulnerability score decrease of 0.18 and 0.2, respectively (Figure S3).

Regardless of the clustering algorithm, three clusters were determined to be the optimal fit for our data by their silhouette coefficients (GMM = 0.43, k-means = 0.457). Slight differences in total number of species per group were observed between the various models (Table S5). The strength of the GMM over k-means clustering is evident when observing the clustering pattern and variation in productivity, susceptibility, and vulnerability between the two methods (Figure S4). When comparing Mclust GMM models, the log-likelihood and BIC values indicated that the “VVI” clustering method is the best fit for our data (Table S5; BIC = 332.738, Log-likelihood = 221.9). This resulted in three vulnerability clusters that have significantly different centroids (ANOVA, p<0.01).

Each cluster represents three distinct vulnerability risk groups shaped by the range of productivity and susceptibility scores within that group. The first group is considered least vulnerable due to their characteristic high productivity and low susceptibility scores (HPLS), followed by a moderately vulnerable group (MPMS), and finally, the vulnerable group is defined by low productivity and high susceptibility (LPHS) (Figure 6). The GMM model shows the HPLS cluster contains 127 species, the MPMS cluster contains 93 species, and the LPHS cluster contains 38 species. The range of vulnerability scores across the clusters are: 0.19-0.73 for HPLS, 0.744-1.11 for MPMS, and 0.87-1.82 for LPHS, the most vulnerable group. The HPLS and MPMS clusters of the GMM model represent fish species that are low priority for further assessment, accounting for 85% of species evaluated. The remaining 38 species (15%) are ranked as vulnerable and are those in need of assessment and management.

**Figure 6:**
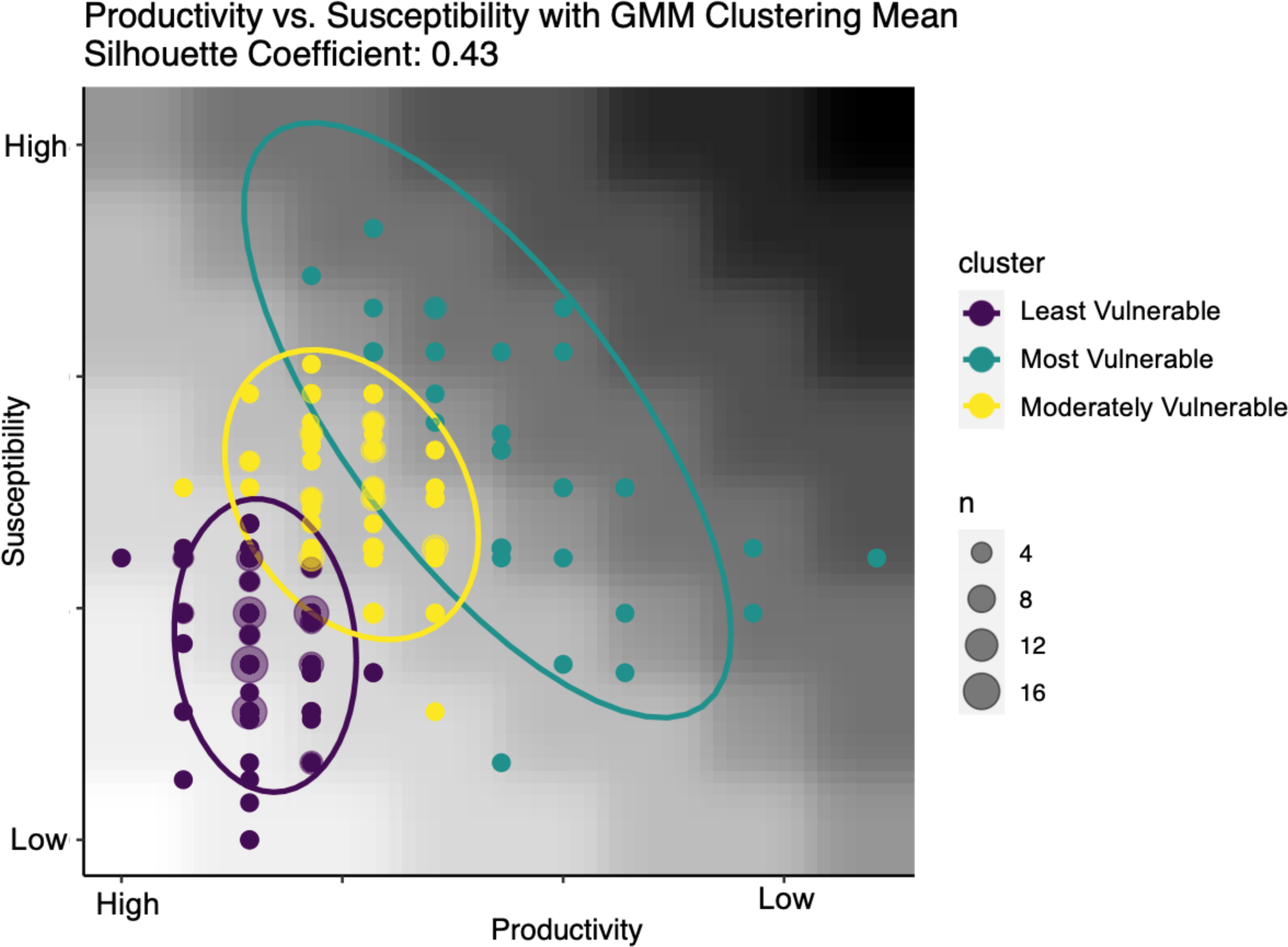
Productivity-Susceptibility Analysis with Gaussian Mixture Model Clustering designating vulnerability categories for all species assessed (n=258). This is a semi-supervised model (silhouette coefficient = 0.43) for grouping individual species into three vulnerability clusters based on productivity, susceptibility, and vulnerability scores. Color gradient represents vulnerability scores with low to high vulnerability scores corresponding with light to dark gradation diagonally from the origin. The size of data points indicate number of individuals with that vulnerability score (see legend).

**Figure 7:**
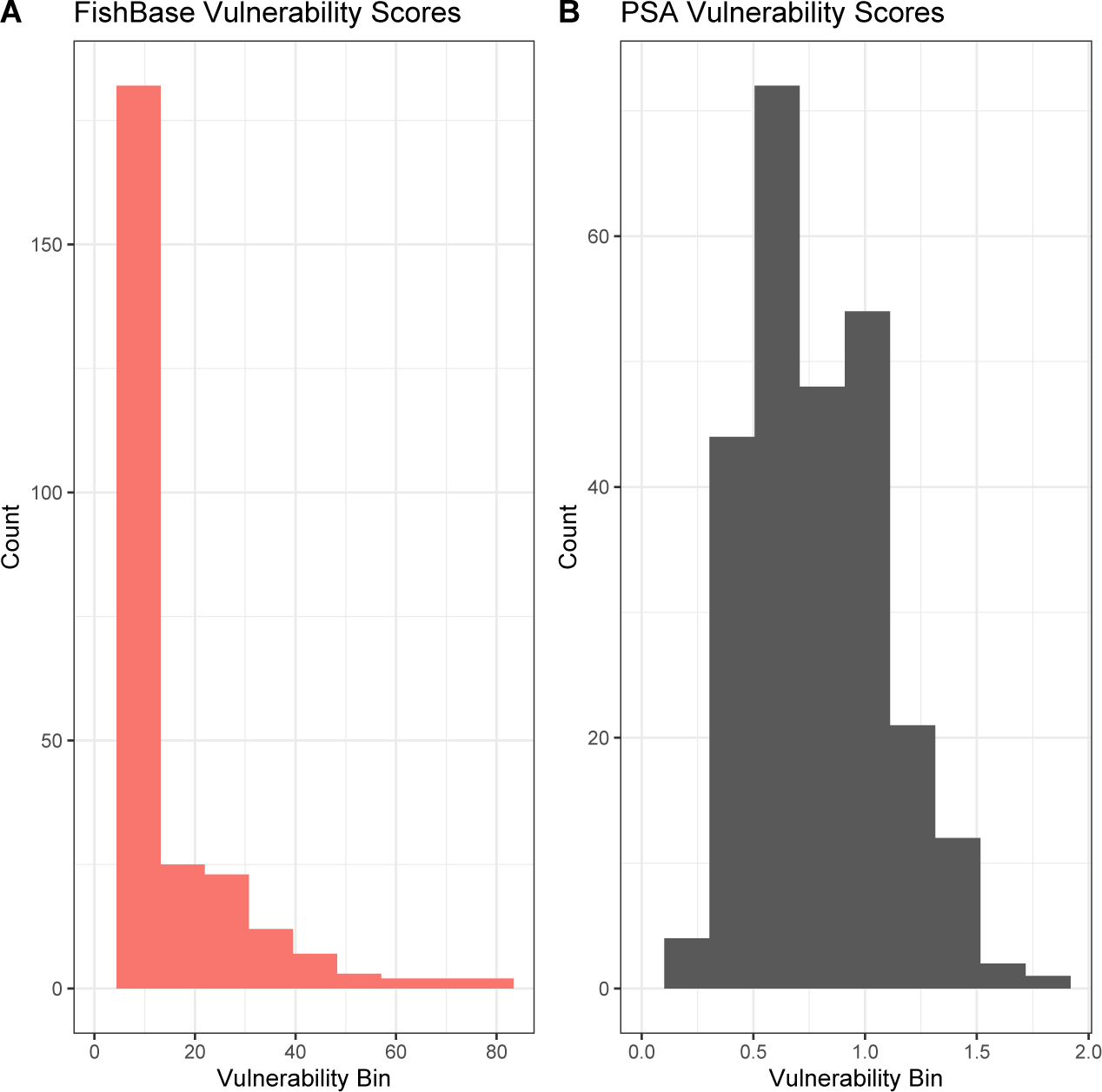
Histogram of vulnerability score distribution (n=10 data bins) from FishBase (A) and the present PSA (B) for all species assessed (n=258). A pair-wise comparison of the FishBase vulnerability data and this PSA show no strong correlation or linear relationship between the two scores for the top 258 fish in the trade (Pearson’s correlation = 0.42, R^2^ = 0.176). The FishBase model (Cheung et al., 2005) is much less sensitive to species-specific characteristics for marine aquarium fish, as 173 species are assessed with a vulnerability of “10”, the lowest possible score in that model. Observing the spread of vulnerability scores between the two models, it is evident the FishBase vulnerability scores are severely left skewed whereas the 2024 PSA scores show a normal distribution when breaking the data into 10 equal vulnerability bins (Figure 7).

The IUCN Red List is widely used as a tool for conservation actions and priorities (IUCN 2022). A majority of the top 258 MAT species are classified as non-threatened in the Red List, with 239 *Least Concern* species and one *Near Threatened* species, *Chiloscyllium punctatum* (IUCN 2022). Three species, namely, *Amblyglyphidodon ternatensis*, *Gobiodon axillaris* and *Chrysiptera hemicyanea* are *Vulnerable* in the Red List due to declining coral cover globally (IUCN 2002). Only one MAT species, *Pterapogon kauderni*, is *Endangered* in the Red List due to its small area of occupancy, severe fragmentation, and continuing decline due to the aquarium trade (Allen *et al*. 2007). We note the significant aquaculture production of *Pterapogon kauderni* (Table 2, Rhyne et al. 2017).

When IUCN Red List results are compared to the PSA, key differences can be noted including the number of factors used, the PSA’s application for data-limited species, and its ability to measure the threat posed by the MAT on species populations. The Red List process uses five criteria to determine a species’ probability of extinction compared to the PSA method that has 11 Productivity and Susceptibility factors for scoring vulnerability to overfishing. The PSA for the 258 MAT species included nine that were considered IUCN *Data Deficient* and five species that have not yet been evaluated in the Red List (IUCN 2022). In the PSA results, the top 10 most vulnerable species in the MAT included *Chiloscyllium punctatum* and *Pterapogon kauderni,* which have *Near Threatened* and *Endangered* conservation status in the IUCN Red List, respectively. The remaining 80% of the top 10 most vulnerable species in the PSA were classified as *Least Concern* in the Red List. Both methods showed that damselfishes (Family: Pomacentridae) are least vulnerable among the top 258 species assessed. Among the species analyzed in the PSA, the 10 species with the lowest vulnerability scores are classified as *Least Concern*, except for *Amblyglyphidodon ternatensis*. This species is categorized as *Vulnerable*, primarily due to the global decline in coral cover (IUCN 2022).

In total, there is 26% overlap between FishBase and this PSA vulnerability index across all species. Comparing the top 15 most vulnerable species ranked by each model, *Chiloscyllium punctatum, Diodon hystrix, Echidna nebulosa,* and *Rhinomuraena quaesita* are the only overlapping species. Comparing seven key species in the trade allows for a further examination of agreement and divergence between these two models (Table 3). The models severely disagreed on the vulnerability score for four *(Pterapogon kaudernii, Paracanthurus hepatus, Dascyllus aruanus, Pomacanthus imperator*) of these seven well-studied species. The damselfish, *Dascyllus aruanus* has opposite vulnerability scores comparing the PSA and FishBase vulnerability scores, respectively (V=0.67, 26) where the main distinction here is the Bangaii cardinalfish (*Pterapogon kaudernii)* scores lower in fishbase vulnerability than the damselfish, while being in the PSA top 10 highest scoring vulnerability species (V=1.52, 19) (Table 3). There is agreement on *Chromis viridis*, and the aforementioned *Echidna nebulosa*.

**Table 3.** Comparison of seven key species vulnerability scores and corresponding vulnerability categories between our PSA and the FishBase Vulnerability model. Species are ordered by low to high FishBase Vulnerability Score.

| Species | PSA<br>Vulnerability | GMM<br>Cluster<br>Vulnerability | FishBase<br>Vulnerability<br>Score | FishBase<br>Vulnerability<br>Category |
| --- | --- | --- | --- | --- |
| *Chromis viridis* | 0.67 | Least | 10 | Low |
| *Sphaeramia*<br>*nematoptera* | 1.2 | Moderately | 10 | Low |
| *Pterapogon kauderni* | 1.52 | Most | 19 | Low |
| *Paracanthurus hepatus* | 1.35 | Moderately | 21 | Low |
| *Dascyllus aruanus* | 0.67 | Least | 26 | Low to moderate |
| *Echidna nebulosa* | 1.56 | Most | 60 | High |
| *Pomacanthus imperator* | 1.02 | Moderately | 68 | High to very high |

## 4. Discussion

We present a PSA tuned for data-deficient species as a robust and accessible tool to predict the vulnerability of fish in the MAT from a holistic and global perspective. This will allow stakeholders to prioritize fish species categorized as vulnerable due to the direct impacts of the marine aquarium fisher, for further study and management consideration. The goal of the PSA is to provide a list of species that are at risk of overexploitation by the trade, specifically identifying the factors that are driving their high vulnerability to inform appropriate management. This PSA represents an expanded species assessment from the framework developed by Baillargeon *et al*. 2020, and increased the number of species considered from 32 to 258, representing the top imported MAT species into the US. It is an adjusted methodology to handle more data-deficient species and serves as a predictive tool to estimate the relative vulnerability of a new fish species entering or becoming more popular in the trade. This framework also identifies fish that should continue to be traded in high volumes or should be the focus of aquaculture production. This PSA acknowledges that local assessment, including site-specific environmental and cultural indicators, should be executed when data are available to provide the most accurate risk assessment and inform strategic management plans.

Sustainable species in the MAT can be defined as those of low susceptibility and high productivity, where a species at risk of overharvesting would display high susceptibility and/or low productivity. This PSA framework does not serve to classify fish into explicit and fixed sustainability categories, but rather a tool to accurately identify species that have a host of characteristics making them resilient or vulnerable to fishing in the MAT. The combination of both where a fish falls on the PSA x-y graph, and in which vulnerability cluster, determines the type and urgency of management plans, respectively. Generally, a PSA assessment can be read based on how far from the origin a species falls. For instance, the top left quadrant indicates a highly susceptible but highly productive species where management would need to be focused on gear limitations as their high productivity will buffer effects of increased fishing pressure. In contrast, the top right quadrant represents a fish with high susceptibility and low productivity, where a greater management focus on protecting broodstock and limiting fishing would be required given the biological demands of that species for them to be sustainable in the trade. A prime example of this is *Echidna nebulosa* (V=1.56) and *Exallias brevis* (V=1.44), which are both within the top 100 species in trade and are within the top 10 highest vulnerability scores for this PSA, consequently in the most vulnerable category. These two species have opposing productivity and susceptibility factors driving their high vulnerability, resulting in different management implications, yet both fall within the most vulnerable cluster.

The majority of fish evaluated by Baillargeon *et al*. (2020) had low susceptibility and high productivity scores, falling in the ‘sustainable’ category, a product of the most traded species being highly fecund with short lifespans (average p = 2.28). Even though we increased our sample size by nearly an order of magnitude, we see similar values in the present analysis (average p = 2.53). However, the vulnerability cluster is dependent on those species included in the analysis and is a relative, not absolute, value. The 2020 model showed 93.3% of species evaluated were least vulnerable (HPLS) or moderately vulnerable (MPMS), while this revised model with an expanded dataset declined to 85%. The species in this analysis which demonstrate high productivity and low susceptibility, are the best wild-caught species to be harvested at significant volumes for the MAT (V=0.19-0.73 for HPLS, and V=0.744-1.11 for MPMS).

In this PSA, productivity and susceptibility scores fell within a narrow range (1.29–2.86), including multiple groups of overlapping points within a single cluster. When expanding the PSA from 32 to 258 species, the general distribution of points mirrors the pattern seen in *Baillargeon et al.* 2020, with 3 distinct ellipse clusters representing the optimal number of vulnerability categories. A key difference in the expanded dataset was the distinct formation of vertical bands of points, indicating large groups of species with the same productivity score. Susceptibility scores are responsible for vertically separating groups of species with similar productivity values, indicative of shared life-history traits, into varying vulnerability score ranges. Although susceptibility was built into the model using a geometric logarithmic mean to avoid heavily skewing the vulnerability score, it remains the most influential factor in determining which vulnerability cluster a species will fall in. For instance, 26 species, or ∼10% of all species assessed in this analysis scored 2.29 for productivity, which is slightly below the median productivity score of 2.57. Of those species, nine fell in the most vulnerable category and 16 fell in the moderately vulnerable cluster. The distinction between the vulnerability cluster directly corresponds to when susceptibility scores are above or below a 1.9 score. With the moderately vulnerable cluster ranging from 1.3-1.8, and most vulnerable ranging from 1.9-2.15 for susceptibility score. For those scoring higher in susceptibility, shifting from one vulnerability cluster to the next indicates a clear change in how a species would be monitored and managed for a given productivity band.

Several species vulnerability scores and rank in trade exemplify why this risk assessment is a useful tool to assess current sustainability and prevent future declines due to fishing activity. In the least vulnerable cluster, *Chrysiptera unimaculata* (V = 0.19) comes close to the lowest possible score, but at the same time, is not well represented in the trade; however, PSA predicts that this species would respond well to increased demand from consumers (rank 190 of 258). *Chiloscyllium punctatum* (V = 1.82 where maximum V=2.82) is the highest scoring species on this list but is also not well represented in trade (252 of the 258 species). In this case, due to their low productivity a consideration of a quota limit may be a next management step to prevent future declines with changes in demand or fishing effort. However, *P. kauderni* is third most vulnerable and is 9th in the trade. This vulnerability of wild populations led the US federal government to initiate a 4D ruling, unfortunately ignoring the significant numbers of this species being produced in aquaculture in Thailand (Rhyne et al. 2017). The PSA framework enables managers to suggest minimally disruptive management methods to decrease risk based on individual productivity and susceptibility scores to avoid implementing broad, uniform management policies that are not tailored to the fishery.

Within this dataset, there were numerous species with limited or unavailable data entirely. This was most often positively correlated with volume in trade. In total, 25% of families comprise a majority of the trade which is why this study took a genus and family level estimation approach for data-deficient factors, as well as identifying trends that hold true at the family and genus level. Half of the productivity factors (Pelagic larval duration, fecundity, and breeding strategy) were data deficient. A step-wise system of estimating data deficient factors by nearest common relative minimizes errors in factor score estimation, an improvement from exclusively family level generalizations. Dynamic yet data-rich susceptibility factors have drastically improved the accuracy of the vulnerability score, while scaling down productivity factors have reduced error propagation.

This improved PSA methodology has two advantages over the IUCN Red List process, making it suitable for determining appropriate management, research, and conservation actions for MAT species. First, this PSA method is best matched to assess the vulnerability of wild-caught fishes in the MAT due to the specialized factor selection to match both the unique life histories and measurable fishing activity for this data-limited system. We were able to calculate a vulnerability score and rank MAT species despite data limitations, including species that were *Data Deficient* in the Red List. Second, this PSA method provides a ranking for a species based on their vulnerability to overfishing in the MAT. Although crucial, the direct impact of the MAT on local coral reef ecosystems is often unavailable (Ochavillo *et al*. 2004). Within the list of *Least Concern* species in the Red List, further prioritization through the PSA was possible in light of the species’ vulnerability in the MAT.

When looking at the types of threats considered by each method, the Red List evaluates the probability of extinction of a species with consideration of all potential threats to its global population. In contrast, the PSA provides a vulnerability score associated with a specific threat, overfishing due to the global trade. A resulting rank for each species assessed in this PSA provides a tool for prioritizing species for research and conservation inclusive of threats from the MAT itself, which is not achieved automatically after examining the conservation status of these species in the IUCN Red List. A threatened category in the IUCN Red List does not necessarily equate to high vulnerability to overfishing in the MAT, exemplified by our results. Except for the *Endangered* cardinalfish, *Pterapogon kauderni*, the three other MAT species were placed in the *Vulnerable* category due to declining coral cover associated with climate change on the Red List. Our improved PSA methodology and resulting ranking is apt for resource managers, policy-makers and researchers looking for information on species vulnerable to the MAT.

Similarly, the nuance of the PSA assessment can be confirmed when comparing our results to that of FishBase Vulnerability outputs (Froese and Pauly, 2023), an automated tool that requires little to no information to be inputted on the user end to obtain the vulnerability score of nearly any fish. The FishBase model is reliant almost entirely on estimated life history parameters rooted in the von-bertalanffy growth equation (Cheung et al., 2005). For a group of diverse, data-deficient species like those in the MAT, this results in either a homogenization of all fish without taking into account any susceptibility factors; or, the fish that do score higher are inflated based on one growth characteristic. Most notably, the Bangaii Cardinal fish (*Pterapogon kaudernii*), the only species demarcated as *Endangered* under the US Endangered Species Act and IUCN Red List, is ranked as low vulnerability (V=19) under the FishBase model. This scoring is directly contradicted by the universal agreement that this fish is endangered due to the wild-caught aquarium fishery, given its unique life history traits of mouth brooding few young in an endemic range while being in high demand due to their appeal to hobbyists, thus making intensive trade unsustainable for this species. Even the fast growing, very fecund damselfish *Dascyllus aruanus* (V=26), scores higher than a known endangered species which is in direct opposition to the known, well-documented life histories and fishing records of both these fish.

The only consistent alignment between the two frameworks is for data-rich fish that behave as traditional food fish in terms of both growth pattern and susceptibility to fishing pressure, like the Snowflake moray eel (*Echidna nebulosa,* V=60). To further demonstrate this point, a popular angelfish, *P. imperator*, has consistently fallen into the moderate risk category across multiple PSA frameworks (Dee et al., 2019; Okewma et al., 2016, Baillargeon et al., 2020). Yet according to the FishBase model it scores as highly vulnerable (V=68) across, in line with the dogfish (*Squalus acanthias*) score (V=68) showing a huge discrepancy in scoring for marine aquarium fish under this framework. Therefore, a model that is not tailored both in what factors it assesses and the mathematical framework (data binning and scoring), data-deficient estimation, and value traits toward aquarium fish instead of food fish fails to be a useful tool to assess and prioritize species at risk.

The cumulative results of our expanded PSA show that the majority of fish in the trade can be considered sustainable. Key traits of a resilient fish in the marine aquarium fishery: Small maximum size, broadcast spawner or high parental investment demersal spawner with high fecundity, wide habitat specificity and large geographic range, are only harvested as juveniles, and are considered easy to care for in a home aquarium. These are often factors that can be assessed on a species or genus level for species in the trade, in lieu of traditional fishery data which is largely unavailable or lacking in granularity. For example, the combination of MAT specific productivity factors replaces the reliance on growth metrics derived from length-weight relationships or spawning stock biomass surveys which are commonly drawn upon to assess the status of a fished stock. If a fish does not meet these exact criteria that does not equate to it being highly vulnerable to overfishing. Instead, the PSA analyzes how well its life history characteristics (productivity factors) are in balance with existing fishing pressure (susceptibility factors). Managers can then read the PSA graph (Figure 6) as four quadrants that require unique management based on if risk is being driven by productivity or susceptibility factors and can tailor management plans to mitigate risk based on this.

Given that this data-limited system operates in some of the most biodiverse coral reefs in the Indo-pacific, which are already facing an onslaught of environmental challenges, there is a need to ensure the marine aquarium industry is providing a positive net benefit to the coral reef socio-ecological system inclusive of reef and community health. The PSA is the best suited data-limited assessment tool to identify the level and type of risk that fishing poses to current and future species that are heavily traded. Its flexible yet robust framework incorporates 11 data points, far more than either FishBase or IUCN, into a single suite of vulnerability indicators (productivity, susceptibility, vulnerability score and cluster) that enable accurate and simple prioritization of species for management.

## 5. Conclusions

We present a robust and accessible risk assessment framework that can be adapted to a diverse range of marine aquarium fish with varied data availability, an essential tool for management bodies such as the IUCN or CITES as the trade continues to draw scrutiny over its sustainability. A key benefit of implementing a PSA is the ability to modify its base framework based on locality of assessment, number of species assessed, and type of fishery. The updates made to the previously published framework for MAT (Baillargeon *et al*. 2020) ensure that species’ vulnerability calculations avoid overestimation of scores through setting more realistic criteria for scoring, elimination of extremely data-limited factors, and productivity-scaling (i.e. volume in trade). The availability of sustainability information for the most popular MAT species will lead to more informed decisions among stakeholders such as fishers, retailers, and consumers when choosing species to harvest, sell, and purchase. A further exploration of this methodology would be to incorporate site-specific environmental and biological covariates alongside economic and cultural indicators across a subset of localized assessments, to strengthen the predictive ability of the model and better aid in developing management plans that are feasible and appropriate based on PSA results.

In summary, this PSA provides the most comprehensive methods to quantitatively assess the sustainability of fish in the MAT, even in cases where data is limited or unavailable for one or more factors. It serves as a useful tool in strengthening other, more qualitative species evaluations. We recommend that the 38 fish in the most vulnerable cluster are prioritized for independent fishery assessment at a national and regional scale to best understand the threat the marine aquarium trade poses to their populations, and how to implement management policies to prevent declines. Further, these species are of high priority for supplementing wild-caught with aquacultured fish. The PSA method can be used in data-limited situations to support IUCN Red List Assessments. The potential impact of this extensive list of vulnerability scores for the top 258 species traded should be viewed as a powerful tool for national and international regulatory bodies, such as CITES or IUCN, to adapt into their risk assessment methodologies when robust datasets are frequently unavailable.

## Acknowledgements

We thank Kent Carpenter, Gina Ralph, and Christi Linardich of the Marine Biodiversity Unit at ODU for technical support and assistance in accessing the IUCN Red List Assessment Database. We also thank and acknowledge feedback from Josie South on the initial review of this manuscript.

## Data Availability

The data that support the findings of this study will be freely available and open-access in the supplementary materials. Volume in trade data can be accessed at aquariumtradedata.org. Supplemental material datasets can be accessed at <u>10.5281/zenodo.10970348</u>.

## Conflict of Interests

Gabrielle A. Baillargeon is funded by UKRI under a NERC Panorama Doctoral Training Program scholarship at the University of Leeds. CASE partners on this scholarship include: Ornamental Aquatic Trade Association (OATA), Center for Environment, Fisheries, and Aquaculture Science (Cefas), and Mars Inc. The data and statements made herein are exclusively the authors and do not necessarily reflect that of funding partners.

Jemelyn Grace Baldisimo (JGB) collaborated on this study as part of her Virginia Sea Grant Graduate Student Fellowship. JGB is a member of the Philippine Aquatic Red List Marine Ornamentals Committee and IUCN Species Survival Commission. The views expressed in this publication do not necessarily reflect those of IUCN.

The designation of geographical entities in this paper, and the presentation of the material, do not imply the expression of any opinion whatsoever on the part of IUCN concerning the legal status of any country, territory, or area, or of its authorities, or concerning the delimitation of its frontiers or boundaries.

Michael Tlusty advises governments on improving wildlife trade data.

Andrew L. Rhyne: Advises public aquaria on the sustainability of living collections and has formal agreements with aquaria concerning aquaculture production of fish species for display. His laboratory specializes in the production of aquarium fish for the trade. Moreover, he has received funding from pet industry groups and non-governmental organizations (NGOs) interested in restoration aquaculture, reduction in cyanide fishing, and animal welfare. Rhyne also co-developed a wildlife trade platform that is currently being tested by a national government as an implementation solution to increase the granularity of wildlife trade data. These relationships could be seen to influence the research presented, although every effort has been made to ensure they have not.

The statements made herein are solely the responsibility of the authors.

