## Supplemental Material for "Evaluating Species at Risk in Data-Limited Fisheries: A Comprehensive Productivity-Susceptibility Analysis of the Most Traded Marine Aquarium Fish"

**Table S1:** Scoring matrix for PSA factors for productivity and susceptibility with corresponding weight and scoring bin based on raw data parameters.

|  | **Productivity Factors** | **Score 1** | **Score 2** | **Score 3** | **Factor Weight** |
| --- | --- | --- | --- | --- | --- |
|  | **Maximum Size (cm)^1^** | >60 | 30-60 | <30 | 2 |
|  | **Trophic Level^1^** | >3.5 | 2.5-3.5 | <2.5 | 1 |
|  | **Breeding Strategy^1^** | Live bearer/mouthbrooder with a fecundity score of 1, or a broadcast/demersal spawner with a fecundity score of 1 | Distinct pairing, demersal spawner | Broadcast Spawner, demersal spawner with high parental care, or mouthbrooder with a fecundity score of 2 or 3 (above 1,000) | 1 |
| **Fecundity (number of eggs produced yearly)^1^** | | <1,000 | 1,000-15,000 | >15,000 | 2 |
| **Pelagic Larval Duration (days)^1^** | | >30 | 14-30 | <14 | 1 |
|  | **Susceptibility Factors** | **Score 3** | **Score 2** | **Score 1** | **Factor Weight** |
|  | **Volume in trade (scaled on productivity score)^1^** | Greater than 3,000 traded and p=0.77-1.9 | Greater than 3,000 traded and p=1.9-2.0  OR  Less than 3,000 and p=0.77-1.9 | <3,000 total volume in trade  OR  p=2-2.3 at any trade volume | 2 |
|  | **Ecological niche + Geographic distribution^2^** | Small geographic range/Narrow habitat specificity | 1. Large geographic range/Narrow habitat specificity  2. Small geographic range/Wide habitat specificity | Large geographic range/Wide habitat specificity | 2 |
|  | **Cyanide Use^1^** | 1. Country of Export: Indonesia, Philippines, Vietnam  2. Family: Chaetodontidae, Pomacanthidae, Acanthuridae | No rating of 2 for this category | Score of 1 if conditions in score 3 do not apply | 1 |
|  | **Encounterability depth (m)^1^** | <10 | 10-30 | >30 | 1 |
|  | **Aquarium Suitability^1^** | “Difficult” care level (grows very large, complex diet, aggressive, high tank mortality rate) | “Moderate” care level | “Easy” care level: (remains at small size, resilient to environmental changes, non-aggressive, less likely to need replacing) | 1 |
|  | **Life cycle stage of harvest**  **1: Recruit, 2: Juvenile, 3: Subadult, 4: Adult** | Harvested at juvenile and adult stages.  (1,2,3,4), (1,2,3), (2,3), (2,3,4) | Harvested only at adult stage  (3), (3,4), (4) | Only harvested at juvenile stage  (1), (1,2), (2) | 2 |

^1^Adapted from Baillargeon, et al (2020)

^2^Adapted from (Rabinowitz, 1981)

**Table S2:  Scoring matrix for scaling of breeding strategy score.**

If a species had a breeding strategy score of 1, but was highly fecund with a value >1,000 (i.e. score of 2 or 3), the breeding strategy score was scaled to a 3. Conversely, species with breeding strategy scores of 3, but low fecundity values <1,000 (i.e. score of 1), breeding strategy scores were changed to 1.

| **Original breeding strategy score** | **Fecundity value** | **Scaled breeding strategy score** |
| --- | --- | --- |
| 1 | > 1000 | 3 |
| 3 | <1000 | 1 |

**Table S3: Scoring matrix for life cycle stage at harvest (LCSH).**

| **Life cycle stage at harvest scoring: Recruit (1), Juvenile (2), Subadult (3), Adult (4)** | | |
| --- | --- | --- |
| **Score 3** | **Score 2** | **Score 1** |
| Harvested at both juvenile and adult stages: (1,2,3,4),(1,2,3),(2,3), (2,3,4) | Harvested only at adult stages: (3), (3,4), (4) | Harvested only at juvenile stages: (1), (1,2), (2) |
| Broodstock and juvenile fish who have not yet reproduced are both removed from fishery | Established broodstock who have made it to adult stages are removed from fishery | Juveniles who may or may not reach maturity are removed from fishery |

**Table S4:  Results of primary (3 factor manipulation) and expanded (single weighted and unweighted) sensitivity analysis.**

|  | **Vulnerability score decrease when shifting from a score of 1 to 3** | **Vulnerability score decrease when shifting from a score of 3 to 1** |
| --- | --- | --- |
| **3 factor manipulation, one factor weighted:** | 0.882 (Fecundity, Breeding strategy, PLD) | 0.734 (Aquarium suitability, encounterability depth, LCSH) |
| **Single weighted factor manipulation:** | 0.364 (Maximum size) | 0.387 (Ecological niche + distribution) |
| **Single unweighted factor manipulation:** | 0.183 (Trophic level) | 0.199 (Aquarium suitability) |

| Number of Species per Cluster | | | |
| --- | --- | --- | --- |
| Cluster Category | **GMM** | **K-Means** | **GMM VVI** |
| Low Vulnerability | 166 (64.3%) | 143 (55.4%) | 127 (49.2%) |
| Moderately Vulnerable | 85 (32.9%) | 83 (32.2%) | 93 (36%) |
| High Vulnerability | 7 (2.7%) | 32 (12.4%) | 38 (14.7%) |
| Vulnerability of Centroid Center Points | | | |
| Low Vulnerability | 0.62 | 0.55 | 0.54 |
| Moderately Vulnerable | 1.07 | 0.99 | 0.92 |
| High Vulnerability | 1.32 | 1.02 | 1.26 |
| Model Validation | | | |
| Silhouette Coefficient | 0.487 | 0.457 | 0.4335 |
| BIC | 1208.6 | 44.64 | 332.738 |
| Log Likelihood | 651.51 | - | 221.90 |

**Table S5:** Comparative table of three different clustering algorithms: Gaussian-Mixture Model (GMM) with equal shape, size, and direction, K-means clustering using kmeans R package, GMM VVI with diagonal clustering and varying volume and shape. Both GMM models were run using Mclust package in R. The number of species per cluster (total % of species assessed) is displayed for each model. The average vulnerability score for each cluster centroid across the three models is shown, along with the silhouette coefficient and BIC value for each model. Log likelihood is only compared between the two GMM models.
